# Sex differences in cocaine self-administration behaviour under Long Access versus Intermittent Access conditions

**DOI:** 10.1101/507343

**Authors:** Hajer Algallal, Florence Allain, Ndeye Aissatou Ndiaye, Anne-Noël Samaha

## Abstract

A widely accepted rodent model to study cocaine addiction involves allowing animals continuous access to drug during long self-administration sessions (‘Long-access’ or LgA). This produces continuously high brain concentrations of drug during each session. This might not model the pharmacokinetics of cocaine use in experienced human users, which are thought to involve intermittently spiking brain cocaine concentrations within and between bouts of use. Intermittent-access (IntA) cocaine self-administration models this spiking pattern in rats. IntA is also particularly effective in increasing incentive motivation for drug. Most IntA studies have been conducted in male rats. Both humans and non-human animals can show sex differences in all phases of the addiction process. We compared cocaine use in female and male rats that self-administered the drug (0.25 mg/kg/injection, i.v.) during 10 daily, 6-h LgA or IntA sessions. Cocaine intake was greatest under LgA, and female LgA rats escalated their intake. However, only IntA rats (both sexes) developed locomotor sensitization to self-administered cocaine and sensitization was greatest in the females. Five and 25 days after the last self-administration session, we quantified incentive motivation for cocaine by measuring breakpoints for the drug (0.083-0.75 mg/kg/injection) under progressive ratio. Breakpoints were similar in IntA and LgA rats. There were no sex differences in breakpoints under LgA. However, under IntA, females reached higher breakpoints for cocaine than males. Thus, LgA might be best suited to study sex differences in cocaine intake, while IntA might be best suited to study sex differences in incentive motivational processes in cocaine addiction.

## Introduction

A widely accepted animal model to study the development of cocaine addiction involves giving animals virtually continuous access to drug for extended sessions (i.e., Long Access sessions; LgA (Ahmed & Koob, 1998). This achieves high and sustained levels of drug intake and brain concentrations of drug (Allain, Bouayad-Gervais & Samaha, 2018; Zimmer, Oleson & Roberts, 2012). However, cocaine intake in human addicts is intermittent, both within and between bouts of intoxication [(Beveridge, Wray, Brewer et al., 2012; Gawin & Kleber, 1986), reviewed in Allain, Minogianis, Roberts et al. (2015)]. This would produce repeated peaks and troughs in blood/brain concentrations of cocaine (Beveridge, Wray, Brewer et al., 2012; Zimmer, Oleson & Roberts, 2012).

Zimmer, Dobrin & Roberts (2011) developed an Intermittent-Access (IntA) self-administration procedure in rats, and it is uniquely effective in producing changes in brain and behaviour that mediate the transition to addiction. In contrast to LgA, where brain concentrations of cocaine remain high for several hours, IntA achieves spikes and troughs in brain concentrations of drug (Allain, Bouayad-Gervais & Samaha, 2018; Zimmer, Dobrin & Roberts, 2011; Zimmer, Oleson & Roberts, 2012). IntA produces much less drug intake than LgA but it more effectively increases incentive motivation for cocaine. IntA rats show greater incentive motivation for cocaine than LgA rats, as measured by behavioral economics measures (Zimmer, Oleson & Roberts, 2012) or breakpoint under a progressive ratio schedule of reinforcement [PR; (Allain, Bouayad-Gervais & Samaha, 2018)]. Increased motivation for cocaine also lasts longer in IntA than in LgA rats (James, Stopper, Zimmer et al., 2018). IntA rats show decreased elasticity of the cocaine demand curve, an increased willingness to respond for cocaine despite electric foot shock, and more cue-induced relapse than generally seen in LgA rats (Kawa, Bentzley & Robinson, 2016). Limited IntA experience is also sufficient to produce such symptoms (Allain & Samaha, 2018; Calipari, Siciliano, Zimmer et al., 2015). Thus, beyond how much drug is taken, the temporal pattern of drug use is decisive in predicting outcome. This is challenging long-held beliefs about what constitutes a good animal model of cocaine addiction (Allain, Bouayad-Gervais & Samaha, 2018; Allain, Minogianis, Roberts et al., 2015; Kawa, Bentzley & Robinson, 2016).

Females and males can respond differently to cocaine (Becker & Hu, 2008). Men are more likely to abuse drugs and to develop addiction (Brady & Randall, 1999). However, women report taking more cocaine (Griffin, Weiss, Mirin et al., 1989), and women can be more vulnerable to relapse to cocaine use after abstinence (Elman, Karlsgodt & Gastfriend, 2001). Women also progress more rapidly from initial cocaine use to entering treatment (Griffin, Weiss, Mirin et al., 1989; McCance-Katz, Carroll & Rounsaville, 1999), they seek treatment more readily (Fiorentine, Anglin, Gil-Rivas et al., 1997) and they enter treatment at a younger age (Kosten, Gawin, Kosten et al., 1993). Biological factors contribute to sex differences in the susceptibility to cocaine addiction. Compared to male rats, female rats are more likely to develop psychomotor sensitization to cocaine (Glick & Hinds, 1984). Psychomotor sensitization is thought to reflect neurobiological changes linked to pathological drug wanting (De Vries, Schoffelmeer, Binnekade et al., 1998; Lorrain, Arnold & Vezina, 2000; Robinson & Berridge, 1993). In agreement, compared to male rats, female rats can acquire intravenous (i.v.) cocaine self-administration sooner (Lynch & Carroll, 1999), they take more cocaine throughout the circadian cycle (Lynch, 2006) and they work harder for cocaine under PR (Lynch, 2006; Roberts, Bennett & Vickers, 1989). Female rats are also more susceptible to cocaine-primed relapse behavior (Lynch & Carroll, 2000), but they could be less susceptible to cue-induced relapse (Fuchs, Evans, Mehta et al., 2005). Specifically under LgA conditions (Roth & Carroll, 2004), female rats take more cocaine than male rats, and they are also more likely to increase their cocaine intake when shifted from short- (1-h) to longer (6-h; LgA) self-administration sessions (Roth & Carroll, 2004).

In a recently published study, it was found that compared to male rats given IntA to cocaine, female IntA rats take more cocaine and show a faster and greater increase in motivation for the dug, as measured by behavioural economics indices (Kawa & Robinson, 2018). Thus, under IntA conditions, females might be more vulnerable to cocaine-induced incentive sensitization, an effect thought to facilitate the transition to addiction (Kawa & Robinson, 2018; Robinson & Berridge, 1993). To assess this further, we determined how IntA versus LgA experience influences sex differences in cocaine consumption patterns, the development of psychomotor sensitization and incentive motivation for the drug.

## Materials and Methods

### Animals

The animal care committee of the Université de Montréal approved all experimental procedures, and these followed the guidelines of the Canadian Council on Animal Care. Adult male (225-250 g) and female (150-175 g) Wistar rats (Charles River Laboratories Saint Constant, QC) were housed individually under a reverse 12h/12h dark/light cycle (lights off at 8:30 am). Experiments were conducted during the dark phase of the rats’ circadian cycle. Water was available ad libitum. Food was restricted to 20 g/day for females and 25 g/day for males. All rats gained weight over days.

### Apparatus

Rats were trained and tested in standard operant cages equipped with two retractable levers (Med Associates, St Albans, VT). Pressing the active lever produced reinforcement (food pellet or intravenous cocaine), pressing the inactive lever had no programmed consequences. At the start of each session, levers were inserted into the cage and the house light was illuminated. During reward delivery and the ensuing timeout period where applicable, both levers were retracted and the light above the active lever was illuminated. The light was then extinguished, and the levers were again inserted into the cage to indicate reward availability. Each cage also contained four horizontally aligned photocell beams to measure locomotion during each self-administration session. Locomotion was computed as photocell beam breaks/min.

### Acquisition of food and cocaine self-administration

As shown in Figure 1 rats were first trained to lever-press for 45-mg banana-flavoured food pellets (grain-based; VWR, Town of Mount-Royal, QC), under a fixed ratio 1 schedule of reinforcement (FR1) with a 20-s timeout period. Sessions lasted 1 h or until 100 pellets were self-administered. Once rats met this acquisition criterion, they were switched to FR3 for at least 2 sessions. When rats reliably self-administered food pellets (∼ 25 pellets/session, on two consecutive sessions), they were and implanted with catheters into the jugular vein (Samaha, Minogianis & Nachar, 2011; Weeks, 1962). Following recovery, rats learned to self-administer cocaine (0.25 mg/kg/injection; Medisca Pharmaceutique, St Laurent, QC; dissolved in 0.9% saline, delivered over 5 s, with a 20-s timeout), as described in Allain, Bouayad-Gervais & Samaha (2018). Next, half of the animals of each sex was given 6-h IntA sessions (IntA; n = 10-11/sex), and the other half was given 6-h LgA sessions (LgA; n = 11-12/sex).

**Figure 1.**
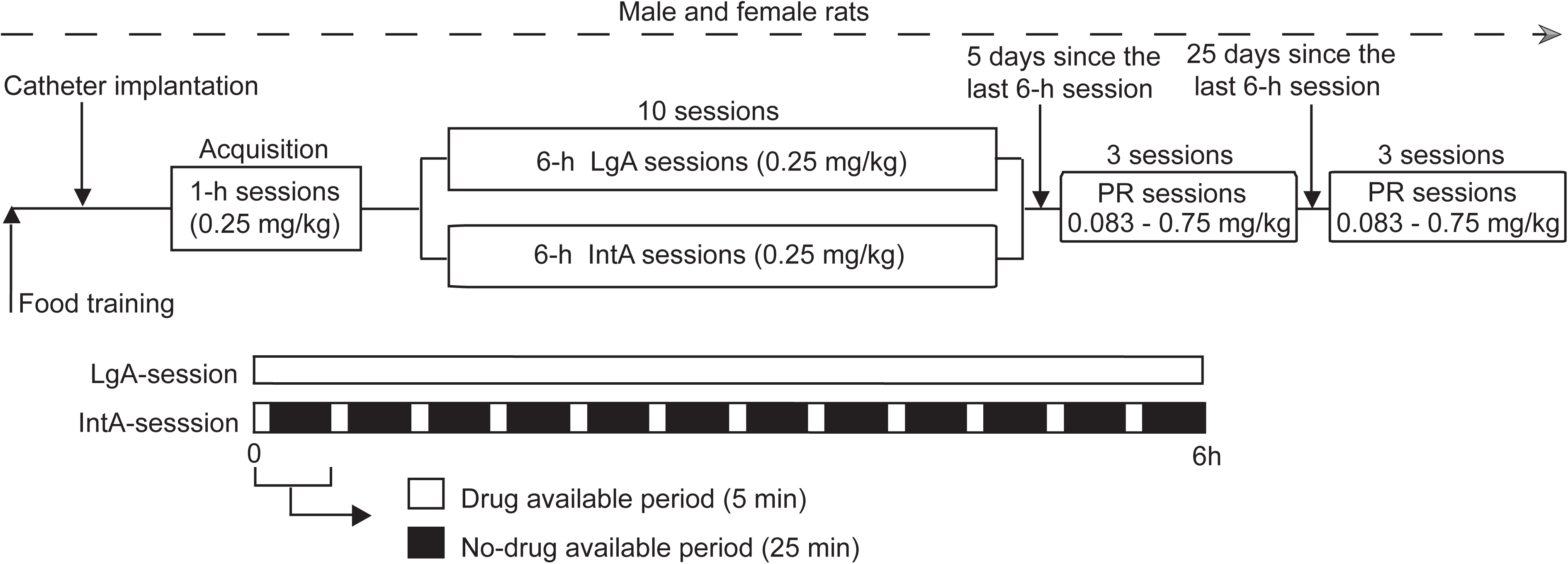
The sequence of experimental events. Male and female rats were trained to press a test lever for banana-flavored food pellets. Subsequently, they were implanted with intravenous catheters and allowed to self-administer cocaine (0.25 mg/kg/injection, delivered intravenously over 5 s) during 1-h sessions. Rats of each sex were then assigned to self-administer cocaine during 6-h Long-Access sessions (LgA-rats) or 6-h Intermittent-Access sessions (IntA-rats). Five and 25 days following the last 6-h self-administration session, incentive motivation for cocaine was assessed by measuring breakpoints for the drug achieved under a progressive ratio schedule of reinforcement (PR).

### IntA and LgA sessions

IntA or LgA sessions (0.25 mg/kg/infusion) were given 1/day, every other day, for 10 sessions. Each IntA-session consisted of twelve 5-min cocaine periods, where drug was available under FR3 without a timeout period (save for each 5-s injection), intercalated with 25-min, no-cocaine periods where levers were retracted (Zimmer, Oleson & Roberts, 2012). During LgA-sessions, cocaine was available continuously under FR3, save for a 20-s timeout following each injection.

### Cocaine self-administration under a progressive ratio schedule of reinforcement (PR)

Five days following the last IntA or LgA session, we assessed breakpoints for cocaine (0.083, 0.5 and 0.75 mg/kg/injection, in counterbalanced order, 1 session/dose) under PR (Minogianis, Levesque & Samaha, 2013; Richardson & Roberts, 1996). A subset of the rats in each group (n = 10-12/sex) was also tested under PR 25 days following the last LgA or IntA session.

### Modeling brain cocaine concentrations

Brain cocaine concentrations (μM) were estimated using self-administration data from the 10^th^ IntA or LgA session, in representative male rats from each group, as in (Allain, Bouayad-Gervais & Samaha, 2018; Zimmer, Dobrin & Roberts, 2011; Zimmer, Oleson & Roberts, 2012). We used a pharmacokinetic model developed and validated in male rats (Pan, Menacherry & Justice, 1991).

### Statistical Analysis

Three-way ANOVA was used to analyze both the number of self-administered injections and locomotion during the 6-h sessions (Sex x Access (LgA or IntA) x Session; Session as a within-subjects variable). Two-way ANOVA was used to analyze cumulative cocaine intake following the ten 6-h sessions. Three-way ANOVA was used to analyze lever pressing behavior during the 6-h sessions (Sex x Lever type x Session; the latter two as within-subjects variables) and breakpoints for cocaine under PR (Sex x Access x Dose; Dose as a within-subjects variable). Significant interaction or main effects were investigated using two-way ANOVA. The statistical significance criterion was *p* ≤ 0.05. Data were analyzed with GraphPad Prism (v. 7.0d) and SPSS (v. 20).

## Results

### Acquisition of food and cocaine self-administration behaviour is similar in male and female rats

Female and male rats acquired reliable food (Figure 2A; *p* > 0.05) and then cocaine (Figure 2B; *p* > 0.05) self-administration behaviour in a similar average number of days. The two sexes also took a similar amount of cocaine during the last two days of acquisition training (Figure 2C; *p* > 0.05).

**Figure 2.**
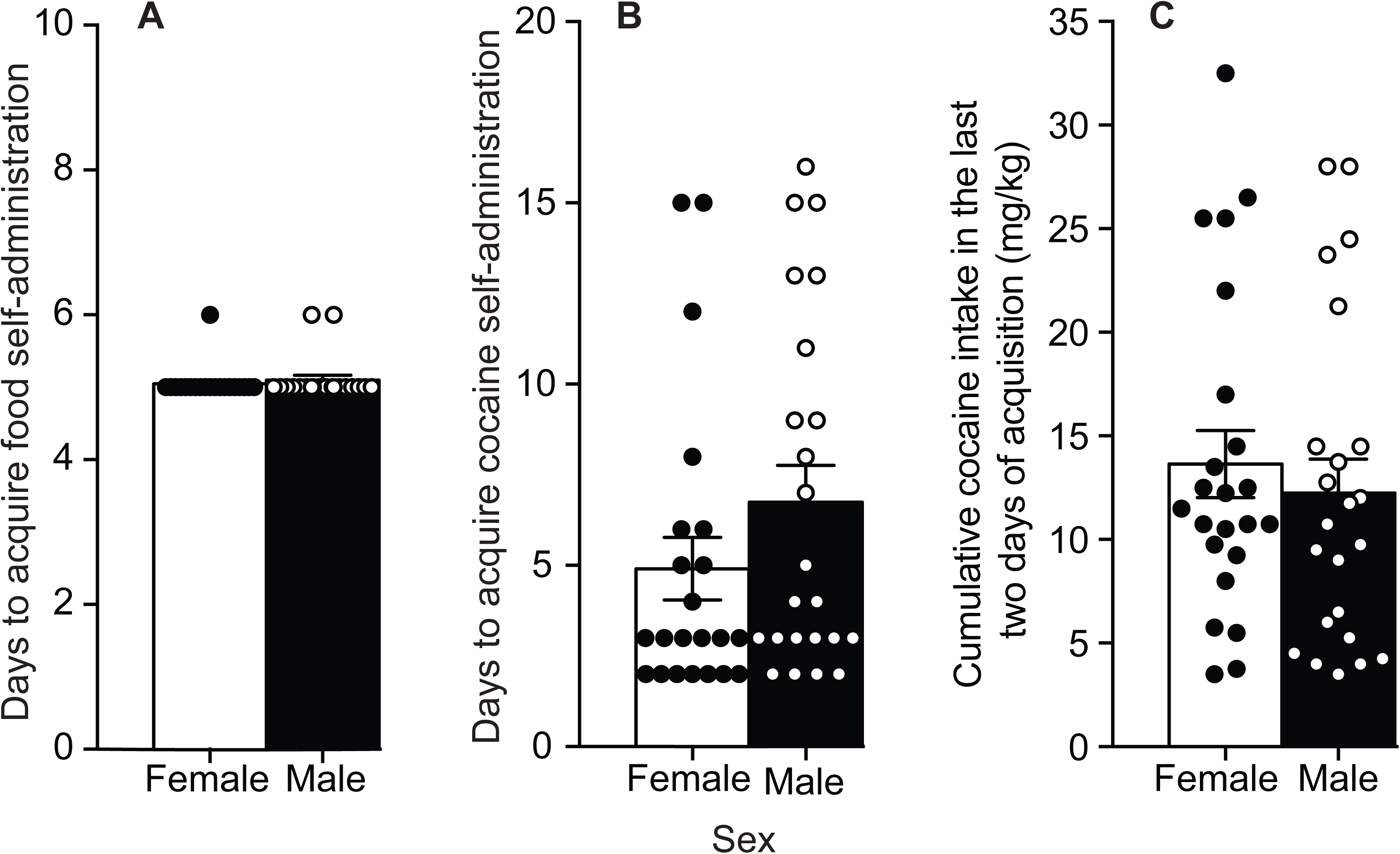
Female and male rats do not differ in the acquisition of food or cocaine self-administration behavior. There were no sex differences in the number of days to acquire (A) food and (B) cocaine self-administration behavior, or in (C) cumulative cocaine intake over the last two days of acquisition training. Data are mean ± SEM. (n = 10 – 12/group).

### Sex differences in the amount of cocaine taken under LgA, but not IntA

Figure 3A shows cocaine intake and estimated brain drug concentrations during the 10^th^ self-administration session in a representative male LgA rat and male IntA rat. Brain cocaine concentrations would be continuously high during a LgA session (Figure 3A, black curve), but would follow a spiking pattern during an IntA session [Figure 3A, grey curve; see also (Allain, Bouayad-Gervais & Samaha, 2018; Zimmer, Oleson & Roberts, 2012).

**Figure 3.**
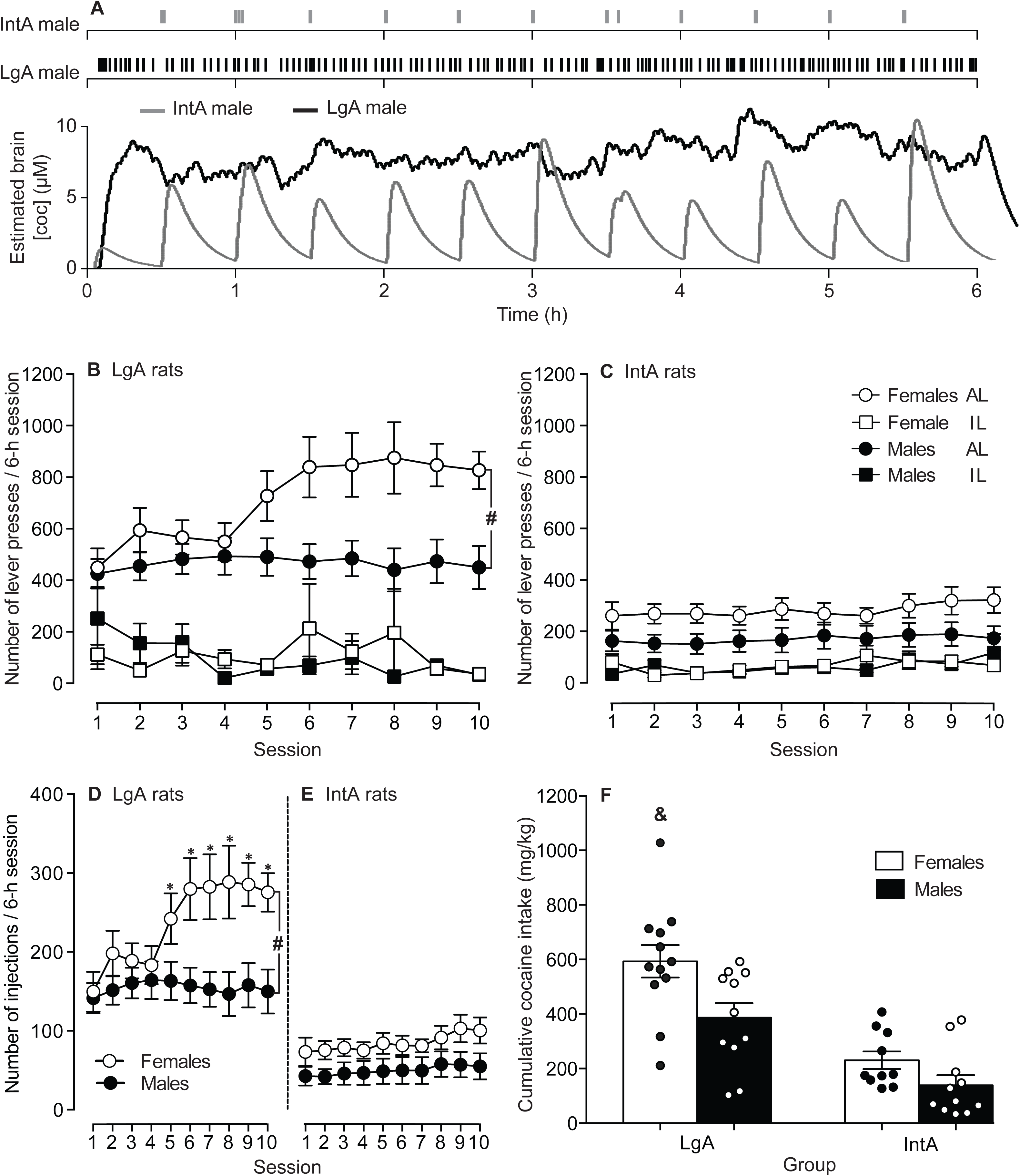
Female rats take significantly more cocaine than male rats under Long-Access (LgA) self-administration conditions, but drug consumption is similar across sexes under Intermittent-Access (IntA). (A) Patterns of cocaine intake and estimated brain cocaine concentrations during the 10^th^ session in representative male rats from the LgA and IntA groups. Under LgA, brain concentrations of cocaine would be continuously elevated. Under IntA, brain drug concentrations would follow a spiking pattern. Under LgA, female rats escalated both (B) their active lever presses and (D) the number of self-administered infusions, but male LgA did not. Under IntA, (C) lever pressing behaviour and (E) the number of self-administered infusions were comparable across the sexes, and did not escalate over time. (F) female LgA rats had the highest levels of cumulative cocaine intake. AL – active lever; IL – inactive lever. ^#^*p* < 0.05. ^*^*p* < 0.0001, vs. the first LgA session in the female rats. &p < 0.05, vs. all other groups. Data are mean ± SEM. n = 10 – 12/group.

Figures 3B-F show cocaine self-administration behaviour over the ten 6-hour sessions. Both LgA and IntA rats reliably discriminated between the two levers, pressing more on the active than on the inactive lever (LgA rats, F_1,21_ = 100.74; Figure 3B; IntA rats, F_1,19_ = 16.06; Figure 3C; All *P*’s ≤ 0.001). Female LgA rats pressed more on the active lever than male LgA rats (Lever type X Sex interaction effect; F_1,21_ = 5.01, *p* = 0.036; Figure 3B), and female LgA rats also significantly escalated their active lever presses (Lever type X Session interaction effect; F_9,21_ = 5.37, *p* < 0.001; Figure 3B). There were no significant sex differences in lever-pressing behaviour in IntA rats (All *P*’s > 0.05). Under LgA, female rats took more cocaine over sessions than male rats (Sex x Session x Access interaction effect; F_9,40_ = 2.96, *p* = 0.002; Figures 3D and E; Main effect of Sex in LgA rats; F_1,21_ = 6.57, *p* = 0.018; Figure 3D). In addition, only LgA females escalated their intake over time (Sex x Session interaction effect; F_9,189_ = 4.25, *p* < 0.001; Figure 3D), such that from the 5^th^ session onwards, they took more cocaine per session than on the 1^st^ session (All *P*’s < 0.05). Under IntA conditions, there was a tendency for female rats to take more cocaine than male rats (Main effect of Sex, F_1,19_ = 3.43, *p* = 0.07; Figure 3E). Cumulative cocaine intake was greater in LgA vs. IntA rats (main effect of Access condition, F_1,40_ = 41.40, *p* < 0.001; Figure 3F), and cumulative intake was greatest in LgA females (All *P’s* < 0.05). Female and male IntA rats took similar amounts of cocaine overall (*p* = 0.07). In summary, LgA rats took more cocaine than IntA rats. In addition, only LgA produced significant sex differences in cocaine intake (0.25 mg/kg/infusion), where females took more cocaine and only females escalated their drug intake over time.

### Sex differences in the pattern of cocaine intake over sessions under both LgA and IntA

As a qualitative measure of the pattern of cocaine intake in the LgA rats, we examined individual cumulative response records on the 1^st^, 5^th^ and 10^th^ sessions in all rats. Figure 4 shows the pattern of cocaine intake during the 1^st^, 5^th^ and 10^th^ LgA sessions in two representative female rats (Figures 4A and B) and two representative male rats (Figures 4C and D). The female rats showed clear escalation of intake from the 1^st^ to the 10^th^ LgA session, while most male rats did not escalate. The female rats also showed two distinct patterns of intake. As illustrated in Figure 4A, some females (4/12) showed bouts of high-frequency cocaine intake followed by brief drug-free periods throughout the LgA session, in particular on the 10^th^ session. As illustrated in Figure 4B, other females (8/12) took closely-spaced injections continuously during the session, and they increased the frequency of cocaine intake over the 10 sessions. In contrast, all males generally had the same pattern of cocaine intake. As seen in Figures 4C-D, the males took took closely-spaced injections almost continuously during the session, and this pattern did not significantly change over sessions.

**Figure 4.**
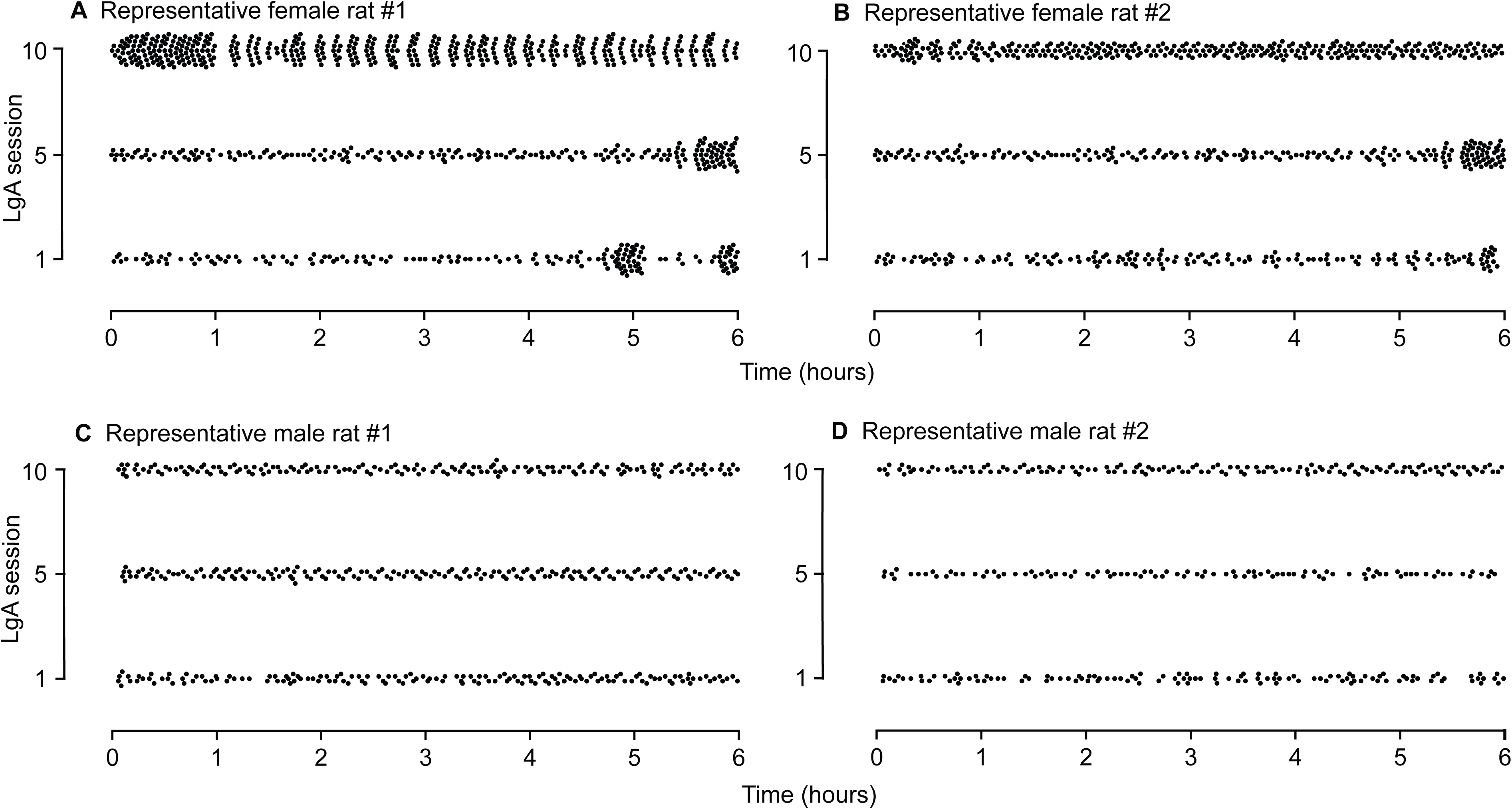
Sex differences in the pattern of cocaine intake under Long-Access (LgA) conditions. Patterns of cocaine intake in representative (A and B) female and (C and D) male LgA rats during the 1^st^, 5^th^ and 10^th^ LgA-sessions. Each point represents one self-administered infusion. Female rats showed two different within-session patterns of cocaine intake. These are illustrated by data from female rat #1 and female rat #2. Female rat #1 consumed cocaine in distinct high-frequency bouts interspersed with brief drug-free periods, in particular on the 10^th^ LgA session. Female rat #2 took closely-spaced injections continuously during each LgA session, with the frequency of intake being highest on the 10^th^ session. The male rats showed less variability in the within-session pattern of intake. As illustrated by male rats #1 and #2, the males took closely-spaced injections throughout each session, with no clear change across sessions. Finally, the female rats significantly escalated their cocaine use over sessions, while the males did not.

We also examined the pattern of drug intake in IntA rats. Male and female IntA rats take most of their cocaine injections within the first 60 s of each drug-available period of the session—taking closely-spaced injections in a burst-like pattern—and this loading effect can sensitize over sessions (Allain, Bouayad-Gervais & Samaha, 2018; Kawa & Robinson, 2018). We examined sex differences in this effect here. Figure 5 shows the pattern of cocaine intake during each 5-min drug period (i.e., 300 seconds) of the 1^st^ (Figures A and D), 5^th^ (Figures 5B-E) and 10^th^ (Figures 5C-F) IntA sessions, in a representative rat from each sex. Looking at these data suggests that both female and male IntA rats took most of their cocaine injections in the first 60 s of each 5-min drug period, and that this loading effect sensitized over sessions in females particularly. To analyse this further, Figures 5G-K show average number of cocaine injections during each minute of the 5-min drug periods, over the ten IntA sessions. Across sessions, both IntA females and males took most of their cocaine in the first minute of each 5-min drug-available period [Time (in 60-s bins) x Session interaction effect; Females; F_36,19_ = 3.56; All *P’*s < 0.05; Males; All *P*’s < 0.05, except 0-60 s vs. 60-120 s, where *p* > 0.05; Figures 5G-K). Over the 10 IntA sessions, female rats also escalated the number of cocaine injections they took in the first minute of each 5-min drug period (*p* = 0.007; Figure 5G), but the male rats did not (*p* > 0.05). To explore this ‘loading’ effect further, we analyzed episodes of burst-like cocaine intake across the 10 IntA-sessions (Figures 5L-P). An episode of burst-like intake was counted when a rat took ≥ 3 injections/60 s (Allain, Bouayad-Gervais & Samaha, 2018; Allain & Samaha, 2018; Belin, Balado, Piazza et al., 2009). Both sexes showed most of their episodes of burst-like intake in the first 60-s of each 5-min cocaine period [Time (in 60-s bins) x IntA session interaction effect; F_36,19_ = 3.58; Main effect of 60-s bin, Females, *F*_1, 18_ = 25.98; Males, *F*_1, 20_ = 6.01; all *P* values < 0.05; Figures 5L-P). However, only females escalated the number of these episodes in the first 60-s bin, such that from the 7^th^ session on, the females showed more of these episodes than on the 1st session (all *P* values < 0.01). In summary, under IntA, both female and male rats took most of their cocaine in the first minute of each 5-min drug period and both sexes showed a burst-like pattern of cocaine intake during this first minute. However, only female rats escalated both their cocaine ‘loading’ effect and their episodes of burst-like intake across sessions.

**Figure 5.**
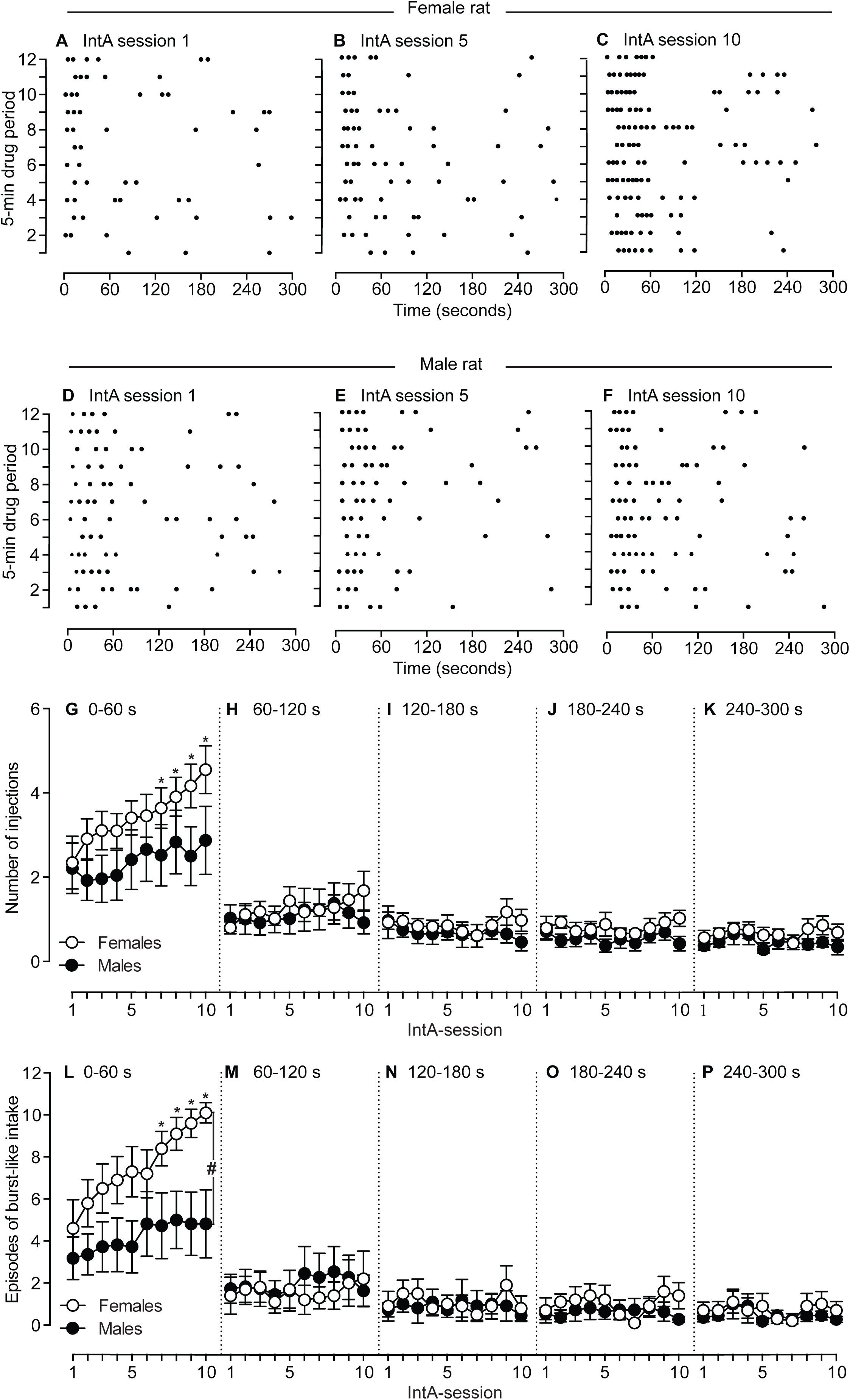
Sex differences in the pattern of cocaine intake under Intermittent-Access (IntA) conditions. Patterns of cocaine intake in a representative (A – C) female and (D – F) male IntA rat during the 1^st^, 5^th^ and 10^th^ IntA-sessions. The Y-axis shows each 5-min cocaine available period of the 6-h IntA session, while the X-axis shows the 5-min drug available period in 60-second (s) bins. Each point represents one self-administered infusion. Both the female and male rat took most of their cocaine injections in the first minute of each 5-min drug period. This is further analysed in (G – K), which show the number of cocaine infusions that females versus males took during (G) the first minute (0 – 60 s), (H) 2^nd^ minute (60 – 120 s), (I) 3^rd^ minute, (120 – 180 s), (J) 4^th^ minute (180 – 240 s) and (K) 5^th^ minute (240 – 300 s) of each cocaine available period across the ten-IntA sessions. (G) Both female and male rats took most of their cocaine infusions in the first minute of each 5-min drug available period, and this ‘loading’ effect significantly sensitized over sessions only in females. (L – P) burst-like events (≥ 3 injections/60 secs) in females versus males during each minute of the 5-min drug available periods, across the ten-IntA sessions. (L) both female and male rats showed most of their burst-like events in the first minute of each 5-min drug available period, and this behavior sensitized significantly over sessions only in female rats. \**p* < 0.05, vs. 1^st^ IntA-session in female rats. ^#^*p* < 0.05. Data are mean ± SEM. n = 10 – 12/group.

### IntA to cocaine promotes locomotor sensitization and this effect is enhanced in female rats

Figure 6A shows locomotor activity in LgA rats during the 10 self-administration sessions. Figure 6B shows locomotor activity in IntA rats, specifically during the 5-min drug-available periods of each session. Cocaine-induced locomotion was greater in IntA rats than in LgA rats (F_1,40_ = 18.83, *p* < 0.0001; Figures 6A-B), and it was also greater in female rats (F_1,40_ = 14.07, *p* = 0.001; Figures 6A-B). Under LgA, there were no significant sex differences in locomotion, and no evidence of locomotor sensitization over sessions. However, under IntA, females showed greater cocaine-induced locomotion across sessions than males (F_1,19_ = 13.11, *p* = 0.005; Figure 6B), and cocaine-induced locomotion increased over sessions in both sexes (F_9,171_= 2.21, *p* = 0.02; Figure 6B). This suggested that IntA rats but not LgA rats, developed cocaine-induced locomotor sensitization. To examine this further, we also analysed locomotor activity during the 25-min no-cocaine periods of the 1^st^ versus 10^th^ IntA session. Compared to male IntA rats, female IntA rats showed greater locomotor activity both on the 1^st^ (F_1,20_ = 131.8, *p* < 0.0001; Figure 6C) and 10^th^ (F_1,20_ = 94.8, *p* < 0.0001; Figure 6C) sessions. In addition, both female and male IntA rats increased their locomotor behavior from the 1^st^ to the 10^th^ sessions (Females, F_1,20_ = 68; Males, F_1,20_ = 4.372; all *P*’s < 0.05; Figure 6C). Thus, both female and male IntA rats developed psychomotor sensitization. To assess potential sex differences in the extent of locomotor sensitization, we compared difference scores in locomotor behavior during the 25-min no-cocaine phases of the 10^th^ versus 1^st^ IntA sessions. This sensitization score was greatest in female rats (*p* < 0.05; Figure 6D). In summary, we did not observe locomotor sensitization in LgA rats. In contrast, both female and male IntA rats showed significant sensitization, and sensitization was more pronounced in females.

**Figure 6.**
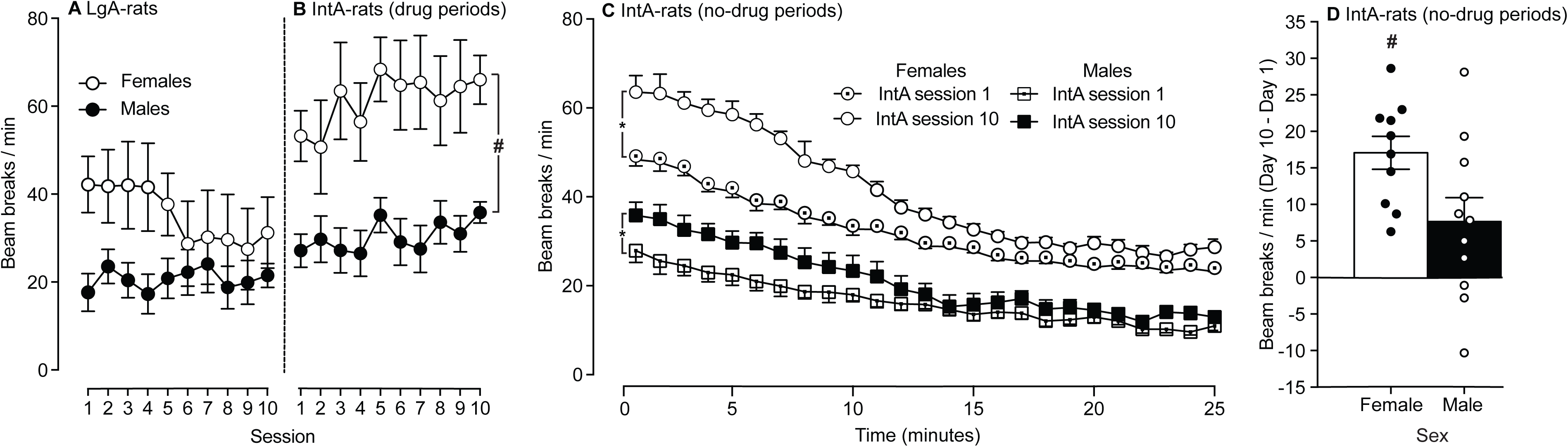
Intermittent-Access (IntA), but not Long-Access (LgA) rats developed robust locomotor sensitization to self-administered cocaine, and sensitization was greatest in female IntA rats. (A) Locomotor activity (measured as beam breaks/min) in female versus male LgA rats over the 10 cocaine self-administration sessions. (B) Locomotor activity in female versus male IntA rats during the drug available periods of each IntA session. The female IntA rats show a greater locomotor response to self-administered cocaine than the male IntA rats. (C) Time course of locomotor activity in female versus male rats during the no drug available periods of the 1^st^ and 10^th^ IntA sessions. In both sexes, cocaine-induced locomotor activity is greater on the 10^th^ versus the 1^st^ IntA session, indicating psychomotor sensitization to self-administered cocaine. (D) The difference in the locomotor score during the no drug available periods of the 10^th^ and 1^st^ IntA sessions in female versus male rats. This score was higher in females, indicating greater psychomotor sensitization. ^#^*p* < 0.001. ^*^*p* < 0.05. ^#^*p* < 0.001. Data are mean ± SEM. n = 10 – 12/group.

### Female rats show more incentive motivation for cocaine than male rats after IntA, but not LgA experience

Five and 25 days after the last 6-h self-administration session, we measured breakpoints for cocaine (0.083-0.75 mg/kg/injection) under PR (Figures 7A-D). At each withdrawal time, female rats reached higher breakpoints for cocaine than male rats, across access conditions (Main effect of Sex; Withdrawal day 5, F_1,40_ = 7.03, Figures 7A-B; Withdrawal day 25, F_1,20_ = 9.01, Figures 7C-D; all *P’s* < 0.02). Cocaine access conditions influenced breakpoints in female rats, but not in male rats. After 5 days of withdrawal, IntA females reached higher breakpoints for cocaine than LgA females (F_1,20_ = 4.73, *p* = 0.04). At each withdrawal time, IntA females also reached higher breakpoints than IntA males (Withdrawal day 5; F_1,19_ = 7.39, Figure 7B; Withdrawal day 25, F_1,9_ = 6.49, Figure 7D; all *P’s* < 0.04). No other comparisons were significant. In summary, after LgA experience, there were no sex differences in incentive motivation for cocaine. After IntA experience, female rats showed greater incentive motivation for cocaine than male rats, both early (5 days) and late (25 days) after cocaine withdrawal.

**Figure 7.**
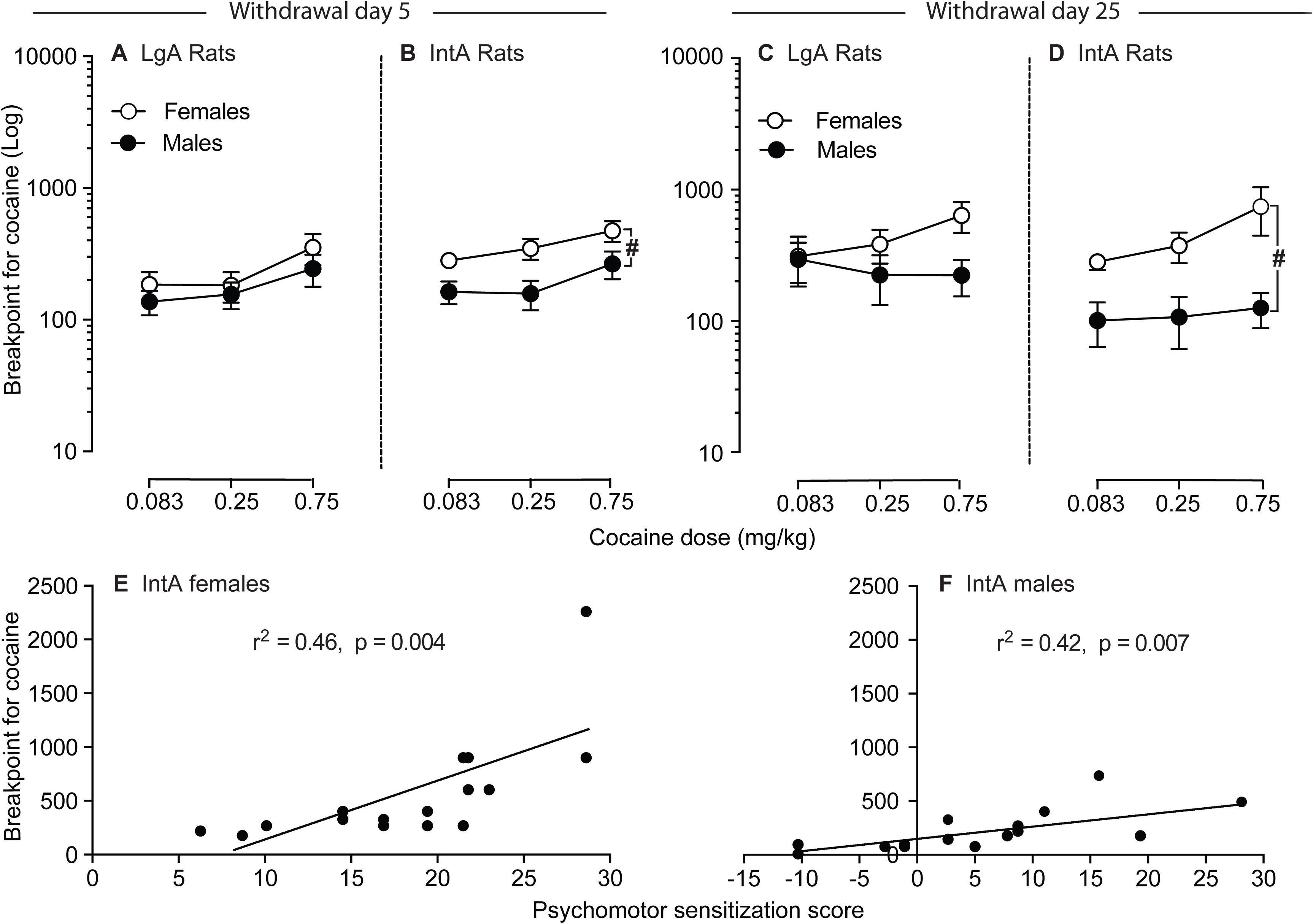
After Intermittent-Access (IntA) experience, breakpoints for cocaine are greater in female versus male rats, but breakpoints are similar across sexes after Long-Access (LgA) experience. Breakpoints for cocaine in (A) LgA and (B) IntA female and male rats, 5 days after the last self-administration session. Breakpoints for cocaine in (C) LgA and (D) IntA female and male rats, 25 days after the last self-administration session. After IntA experience, female rats reached higher breakpoints for cocaine than male rats, at each time after cocaine withdrawal. After LgA experience, there were no sex differences in breakpoints, at any time after cocaine withdrawal. ^#^*p* < 0.01. In both (E) female and (F) male IntA rats, there was a significant positive correlation between the degree of psychomotor sensitization (difference scores in locomotor activity/min during the no-cocaine available periods of the 10^th^ versus 1^st^ IntA sessions) and subsequent breakpoints for cocaine (0.75 mg/kg/infusion). Data are mean ± SEM. n = 10 – 12/group on withdrawal day 5; 5 – 7/group on withdrawal day 25.

## The extent of psychomotor sensitization to cocaine predicts incentive motivation for the drug in female and male IntA rats

We determined whether the extent of psychomotor sensitization (the scores in Figure 6D) predicted breakpoint for cocaine (0.75 mg/kg/injection) in the IntA rats (LgA rats did not show psychomotor sensitization). Because only a subset of the rats was tested under PR 25 days after cocaine withdrawal, we pooled the rats across withdrawal times for this correlational analysis. Figures 7E and F show that higher levels of psychomotor sensitization predicted higher levels of incentive motivation for cocaine, in both sexes (Females, *r*^2^ = 0.46, *p* = 0.004, Figures 7E; Males, *r*^2^ = 0.42, *p* = 0.007, Figures 7F). As seen in Figure 7E, one IntA female had high breakpoint and psychomotor sensitization values. When data from this rat are excluded, there is still a significant positive correlation between these two measures (*r*^2^ = 0.54, *p* = 0.002).

## The amount of cocaine taken in the past significantly predicts incentive motivation for the drug only in male LgA rats

Table 1 shows that there was a significant positive correlation between cumulative cocaine intake (total number of cocaine injections taken over the ten 6-h sessions, multiplied by 0.25 mg/kg/injection) and breakpoints for cocaine (0.75 mg/kg/injection; similar results were obtained using breakpoints for 0.083 or 0.25 mg/kg/infusion cocaine) in male LgA rats only (r^2^ = 0.56, *p* = 0.003). Thus, male LgA rats that took high amounts of drug in the past later showed high incentive motivation for the drug. There was no significant relationship between cumulative cocaine intake and subsequent breakpoints for cocaine in the other groups (All *P*’s > 0.05; Table 1).

**Table 1.** Correlations between breakpoints for cocaine (0.75 mg/kg/infusion) under progressive ratio and past cumulative cocaine intake (mg/kg):

| **Group** | **r<sup>2</sup> value** | **p value** |
| --- | --- | --- |
| Male IntA rats | 0.20 | 0.08 |
| Female IntA rats | 0.01 | 0.80 |
| Male LgA rats | 0.56* | 0.003* |
| Female LgA rats | 0.16 | 0.11 |

\* significant positive correlation between cumulative cocaine intake and breakpoint for the drug.

As mentioned above, during LgA sessions, some female rats took cocaine in distinct, high-frequency bouts. Others did not. The two phenotypes did not produce differences in responding for cocaine under PR (0.083-0.75 mg/kg/infusion; *p* > 0.05; data not shown).

## Discussion

We assessed sex differences in cocaine self-administration behavior in rats given IntA versus LgA experience. Consistent with prior work (Becker & Koob, 2016; Lynch, 2018), females were more vulnerable to the reinforcing, psychomotor sensitizing and incentive motivational effects of cocaine than males. Importantly, drug access conditions (LgA vs. IntA) influenced sex differences in the response to chronic cocaine intake. Sex differences in the sensitivity to cocaine reinforcement were more readily observed with LgA. Sex differences in both locomotor sensitization and incentive motivation for cocaine were more readily observed with IntA. Thus, the LgA procedure was more effective in producing sex differences in the amount of cocaine consummed, while the IntA procedure was more effective in producing sex differences in sensitization to the locomotor activating and incentive motivational properties of the drug [see also (Kawa 2018)].

Female and male animals can differ in the rate at which they learn to self-administer cocaine, but we did not observe this here, consistent with a recent IntA study (Kawa & Robinson, 2018). Using continuous-access procedures, some studies report that female rats acquire cocaine self-administration more readily than male rats (Hu, Crombag, Robinson et al., 2004). Others report the opposite (Caine, Bowen, Yu et al., 2004). The sexes might not differ in acquisition of cocaine self-administration behaviour when test conditions promote rapid acquisition [e.g., higher cocaine doses, food restriction, operant pretraining; (Lynch & Taylor, 2004) and references therein]. Some of these conditions resemble ours. However, fully assessing sex differences in the rate of acquisition of cocaine self-administration behaviour requires testing several cocaine doses and schedules of reinforcement.

Under LgA, female rats self-administered more cocaine than males, and only females escalated their intake. This is consistent with the literature showing that females can take more cocaine and they can also be more likely to escalate intake (Becker & Hu, 2008; Lynch & Carroll, 1999; Roberts, Bennett & Vickers, 1989; Roth & Carroll, 2004). Our male LgA rats did not escalate, similar to other LgA studies using lower cocaine doses [0.25-0.6 mg/kg/infusion; (Mantsch, Yuferov, Mathieu-Kia et al., 2004; Minogianis, Levesque & Samaha, 2013)]. Under IntA, female and male rats took similar amounts of cocaine. However, when given access to higher cocaine doses, for longer, IntA females can take more drug than IntA males (Kawa & Robinson, 2018). This suggests that sex differences in cocaine intake under IntA might require more extensive drug exposure. Beyond the amount of drug taken, taking cocaine in a burst-like pattern is thought to contribute to the development of addiction-like symptoms in rats (Belin, Balado, Piazza et al., 2009; Martin-Garcia, Courtin, Renault et al., 2014). Under IntA, we found that both sexes took most of their cocaine at the beginning of each cocaine-available period, taking closely-spaced injections in a burst-like pattern (≥ 3 injections/60 s) and this loading effect sensitized over sessions only in females. This is generally consistent with prior findings in male (Allain, Bouayad-Gervais & Samaha, 2018; Allain & Samaha, 2018; Kawa, Bentzley & Robinson, 2016) and female (Kawa & Robinson, 2018) IntA rats.

Kawa & Robinson (2018) propose that intermittent cocaine use might more readily produce sensitization-related changes in brain motivation pathways in females, thus accelerating the addiction process, and our findings support this idea. Female and male rats took similar amounts of cocaine under IntA, but females developed more robust sensitization to the psychomotor activating and incentive motivational effects of the drug. Psychomotor sensitization is thought to reflect brain changes that lead to sensitized drug wanting, thereby increasing the risk of addiction. IntA cocaine experience produces psychomotor sensitization [(Allain, Roberts, Levesque et al., 2017; Allain & Samaha, 2018) and present data], and the more an IntA rat is sensitized to cocaine the more it shows incentive motivation for the drug (Allain, Roberts, Levesque et al., 2017). This prior work was done in male rats only. Here we show that in both sexes, IntA cocaine experience promotes locomotor sensitization, and the extent of sensitization predicts incentive motivation for cocaine. We found no evidence of sensitization in the LgA rats. However, we did not assess potential stereotyped movements or psychomotor sensitization after a withdrawal period, both of which can make a difference (Ferrario, Gorny, Crombag et al., 2005). The finding that psychomotor sensitization was greater in female versus male IntA rats also agrees with prior work using intermittent, experimenter-administered drug (Becker & Hu, 2008; Glick & Hinds, 1984; Robinson & Becker, 1986). Thus, our findings bring together the literatures on sex differences in sensitization and drug self-administration and show that female rats might be more susceptible to sensitization-related neuroadaptations following voluntary IntA cocaine use [see also (Kawa & Robinson, 2018)]. This is reminiscent of clinical observations, where women enter treatment following a shorter period of cocaine use (Griffin, Weiss, Mirin et al., 1989; McCance-Katz, Carroll & Rounsaville, 1999), suggesting a more rapid course of addiction. Thus, it is possible that women transition faster to cocaine addiction because they are more vulnerable to sensitization-related neuroplasticity induced by the drug.

Compared to LgA cocaine experience, IntA more effectively produces sensitization of incentive motivation for the drug (Allain, Bouayad-Gervais & Samaha, 2018; Zimmer, Oleson & Roberts, 2012). In agreement with this, IntA female rats showed more incentive motivation for cocaine than LgA female rats here, even though the IntA females had taken 3 times less cocaine in the past. In the males, access condition did not significantly influence incentive motivation for cocaine. This contrasts with previous studies (Allain, Bouayad-Gervais & Samaha, 2018; Zimmer, Oleson & Roberts, 2012). Differences between studies in terms of cocaine doses and number of self-administration sessions could explain the discrepancy. These issues notwithstanding, our findings support the idea that IntA cocaine experience is uniquely effective in increasing incentive motivation for drug. Across sexes, LgA rats took 2 to 3 times more cocaine than IntA rats. Yet, compared to LgA, IntA led to greater (in the females) or similar (in the males) motivation to work for cocaine. These results are consistent with others showing that intermittent ‘spikes’ in brain cocaine concentrations are more effective than high and escalating brain concentrations in producing symptoms relevant to cocaine addiction (Allain, Bouayad-Gervais & Samaha, 2018; James, Stopper, Zimmer et al., 2018; Kawa, Bentzley & Robinson, 2016; Zimmer, Oleson & Roberts, 2012).

There could be a number of explanations for the sex differences we observed. Perhaps more drug gets to the brain in females with each self-administered injection. We did not measure brain cocaine concentrations. However, adult, non-anaesthetized female and male rats have similar cocaine concentrations in both plasma and brain after intraperitoneal administration (Bowman, Vaughan, Walker et al., 1999). In both humans (Mendelson, Mello, Sholar et al., 1999) and non-human primates (Mendelson, Mello & Negus, 1999), i.v. cocaine administration also produces similar blood concentrations of the drug in females and males. In addition, gonadal hormones have effects in the brain that can contribute to addiction (Lynch, 2018). We did not monitor estrous cycle here and estrous phase could influence our outcome measures. For example, females reach higher breakpoints for cocaine when estradiol levels are high (Roberts, Bennett & Vickers, 1989). The current findings, along with recently published work (Kawa & Robinson, 2018) are an initial step in characterizing the effects of IntA cocaine intake in female and male animals. As a first step,’. *inclusion of intact females, without regard to estrous cycle, and intact males is a valid approach to learn about females in neuroscience research*’ (Becker, Prendergast & Liang, 2016). In parallel, sex differences in the response to cocaine are also seen in the absence of gonadal hormones, suggesting that the brain systems that mediate cocaine’s effects could also differ between the sexes (Hu, Crombag, Robinson et al., 2004). Of note, as seen in many preclinical and clinical neuroscience studies (Maney, 2016), our female and male animals also overlapped extensively on all behavioural measures. We illustrate this with individual values in some of the figures. Thus, in considering the amount and pattern of cocaine intake, psychomotor sensitization and incentive motivation for cocaine, there is not one phenotype typical of females and the other typical of males. This suggests that factors in addition to sex contribute to variation in the response to cocaine with IntA or LgA experience.

## Conclusion

Compared to male animals, female animals—including in humans—can be more susceptible to cocaine effects that contribute to addiction. Our findings suggest that LgA procedures could be useful to model sex differences in the positive reinforcing effects of cocaine, as measured by drug consumption under fixed ratio schedules of reinforcement. In parallel, IntA procedures could be better suited to study sex differences in sensitization-related neuroadapatations that lead to increased incentive motivation for cocaine, as measured by appetitive responding for the drug. This has implications for the design of studies examining sex differences in the neurobiological, psychological and behavioral response to cocaine at different stages of the addiction process.

## Acknowledgements

This work was supported by grants to ANS from the Canada Foundation for Innovation (grant number 24326) and the Canadian Institutes of Health Research (grant number 200200). ANS is supported by a salary grant from the Fonds de Recherche du Québec – Santé (grant number 28988).

## Author Contributions

ANS designed the experiments. HA and NAN performed the experiments with guidance from FA. HA and FA analyzed the data. HA and ANS wrote the manuscript with input from FA. All authors critically reviewed the content and approved the final version for publication.

